# Nutritional combinatorial impact on the gut microbiota and plasma short-chain fatty acids levels in the prevention of mammary cancer in Her2/neu estrogen receptor-negative transgenic mice

**DOI:** 10.1101/2020.06.08.139865

**Authors:** Manvi Sharma, Itika Arora, Matthew L. Stoll, Yuanyuan Li, Casey D. Morrow, Stephen Barnes, Taylor F. Berryhill, Shizhao Li, Trygve O. Tollefsbol

**Affiliations:** Department of Biology, University of Alabama at Birmingham, Birmingham, AL, United States; Division of Pediatric Rheumatology, University of Alabama at Birmingham, Birmingham, AL, United States; Department of Obstetrics, Gynecology & Women’s Heath, University of Missouri, Columbia, MO, United States; Department of Surgery, University of Missouri, Columbia, MO, United States; Department of Cell, Developmental & Integrative Biology, University of Alabama at Birmingham, Birmingham, AL, United States; Comprehensive Cancer Center, University of Alabama at Birmingham, Birmingham, AL, United States; Targeted Metabolomics and Proteomics Laboratory, University of Alabama at Birmingham, Birmingham, AL, United States; Department of Pharmacology and Toxicology, University of Alabama at Birmingham, Birmingham, AL, United States; Comprehensive Center for Healthy Aging, University of Alabama at Birmingham, Birmingham, AL, United States; Nutrition Obesity Research Center, University of Alabama at Birmingham, Birmingham, AL, United States; Comprehensive Diabetes Center, University of Alabama at Birmingham, Birmingham, AL, United States

**Keywords:** broccoli sprouts, green tea polyphenols, combination, gut microbiota, diet, breast cancer, Her2/neu mice, tumor, short chain fatty acids, metabolome

## Abstract

Breast cancer is the second leading cause of cancer-related mortality in women. Various nutritional compounds possess anti-carcinogenic properties which may be mediated through their effects on the gut microbiota and its production of short-chain fatty acids (SCFAs) for the prevention of breast cancer. We evaluated the impact of broccoli sprouts (BSp), green tea polyphenols (GTPs) and their combination on the gut microbiota and SCFAs metabolism from the microbiota in Her2/neu transgenic mice that spontaneously develop estrogen receptor-negative [ER (-)] mammary tumors. The mice were grouped based on the dietary treatment: control, BSp, GTPs or their combination from beginning in early life (BE) or life-long from conception (LC). We found that the combination group showed the strongest inhibiting effect on tumor growth volume and a significant increase in tumor latency. BSp treatment was integrally more efficacious than the GTPs group when compared to the control group. There was similar clustering of microbiota of BSp-fed mice with combination-fed mice, and GTPs-fed mice with control-fed mice at pre-tumor and post-tumor in both BE and LC groups. The mice on all dietary treatment groups incurred a significant increase of *Adlercreutzia* genus and S24-7 family in the both BE and LC groups. We found no change in SCFAs levels in the plasma of BSp-fed, GTPs-fed and combination-fed mice of the BE group. Marked changes were observed in the mice of the LC group consisting of significant increases in propionate and isobutyrate in GTPs-fed and combination-fed mice. These studies indicate that nutrients such as BSp and GTPs differentially affect the gut microbial composition in both the BE and LC groups and the key metabolites (SCFAs) levels in the LC group. The findings also suggest that temporal factors related to different time windows of consumption during the life-span can have a promising influence on the gut microbial composition, SCFAs profiles and ER (-) breast cancer prevention.

## Introduction

Breast cancer is a significant health concern worldwide as it is the second most common cause of cancer-related mortality among women. In 2019, about 268,600 new cases of breast cancer were diagnosed, which accounted for 30% of all new cancer cases diagnosed in women living in the United States [1]. Breast cancer has been categorized into five main types: luminal A, luminal B, triple negative/basal like (TNBC), Her2-enriched and normal like, on the basis of hormone receptors (estrogen receptor [ER] and progesterone receptor [PR]), human epidermal growth factor receptor 2 [HER2], and Ki67 [2, 3]. Estrogen receptor-positive [ER (+)] (luminal A/B) breast cancer patients can receive hormone therapy with anti-estrogens and/or aromatase inhibitors, and thereby have better treatment prognosis [4]. By contrast, estrogen receptor-negative [ER (-)] (HER2 and basal-types) breast cancer patients have a poor prognosis and fewer cancer prevention and treatment options due to lack of target-directed approaches and the aggressive nature of this disease [5]. The commonly employed treatment approaches for ER (-) breast cancer patients and metastatic breast cancer patients are surgical, chemotherapy, radiation therapy, and palliative therapy [6]. However, these procedures have an array of short-term or long-term side effects in the patients such as loss of hair, vomiting, skin disorders, fatigue, nausea, anemia, diarrhea, muscle disorder, and nerve diseases [6, 7]. Therefore, there is a need for effective and safe approaches for prevention and treatment of ER (-) breast cancer.

The use of dietary bioactive botanicals is considered as a key alternative approach for prevention, progression and treatment of ER (-) breast cancer due to their efficacy and safe consumption in humans [8]. For example, broccoli sprouts (BSp) and green tea polyphenols (GTPs) have been reported to reduce the incidence of breast cancer [9, 10]. Sulforaphane (SFN) is an isothiocyanate present in cruciferous vegetables such as BSp, kale, bok choy, cauliflower and cabbage and also has chemopreventive/chemotherapeutic effects against numerous types of cancers via epigenetic mechanisms [11–13]. Studies have shown that SFN is a potent inhibitor of histone deacetylase (HDAC), which is an enzyme that modulates epigenetic machinery by removal of an acetyl group from histone residues. Studies have shown that SFN induced G1/S arrest, led to down-regulation of SEI-1 and cyclin D2, increased levels of p21 and p27 and promoted cellular senescence in breast cancer [14, 15]. (-)-Epigallocatechin-3-gallate (EGCG), a major polyphenol in green tea, induces epigenetic modulations such as inhibition of DNA methytransferases (DNMTs) and has numerous anticarcinogenic properties both *in vitro* and *in vivo* against several cancers including breast cancer [16–20]. The anti-tumor mechanisms of GTPs and EGCG involve induction of cell-cycle arrest, mitochondrial-mediated apoptosis, inhibition of IL-6 and induction of tumor necrosis factor-α expression, inhibition of enzymes that regulate the glycolytic process and repression of glucose metabolism [6, 21–23].

Our previous studies have shown that the combination of BSp and GTPs resulted in synergistic inhibition of cellular proliferation, ERα reactivation via regulation of DNMT1 and HDAC1 expression in the ERα (-) breast cancer cell lines MDA-MB-231 and MDA-MB-157, and also resulted in a significant inhibition of tumor development in an ER (-) xenograft mouse model [24]. Additionally, these combined dietary components induced cellular apoptosis and cell cycle arrest in the transformed breast cancer SHR cells (normal human mammary epithelial cells transfected with *SV40*, *hTERT* and *H-Ras* genes), and led to genome-wide epigenetic alterations. This combination treatment administered in a breast cancer xenograft mouse model also resulted in significant inhibition of tumor growth when compared with singly administrated compounds [25]. In addition, our recent study reported that the prenatal or maternal consumption of BSp has more protective effects on tumor development than postnatal or adulthood administration of BSp in SV40 transgenic and Her2/neu transgenic mouse models [26].

The human gastrointestinal tract harbors trillions of microorganisms (≥10^14^) that are reported to be at an approximate ratio of 1:1 with the human cells [27]. The gut microbiota plays an important role in regulation of human metabolic and physiological functions by production of crucial metabolites such as short chain fatty acids (SCFAs) [28, 29]. SCFAs are produced from the fermentation of non-digestible carbohydrates by gut microbiota and the major SCFAs include butyrate, acetate and propionate [30]. These microbial-produced metabolites actively participate in epigenetic modulations in the host cells, that in turn can have profound effects on the inhibition of cancer [31].

Several studies have attempted to investigate the link between dietary compounds, gut microbiota and breast cancer [32, 33]. However, few studies have explored the impact of BSp or GTPs on the gut microbiota in relation to the breast cancer. Some studies have shown that BSp or GTPs can have a significant impact on gut microbial diversity and metabolite production in humans and animals [34–37]. BSp are rich in glucosinolates, which are metabolized by gut microbiota into isothiocyanates. Further, dietary supplementation with broccoli was reported to lead to alterations in cecal microbiota composition, metabolism and intestinal morphology in an inflammatory disease mouse model [38]. A recent study focused on the impact of broccoli ingestion on gut microbiota of C57BL/6 mice found that increased levels of *Clostridiaceae*, *Lachnospiraceae* and *Porphyromonadaceae* and abundance in gut microbiota diversity was associated with broccoli consumption [34]. EGCG can be degraded by microbial enzymes produced in the digestive tract [39]. Previous studies focused on the consumption of GTPs on intestinal microbiota of healthy humans have found a significant decrease in *Clostridium* spp. and an increase in *Bifidobacterium* spp. resulting in significantly high levels of acetate and propionate [40]. Others have demonstrated that supplementation of green tea extract resulted in an increase of *Bifidobacterium* species in calves and humans. *Bifidobacterium* is a beneficial bacterial species that possess prebiotic properties and can lead to improvement in colon environment [41, 42]. A deeper understanding of the gut microbial mechanisms underlying the protective effect of BSp and GTPs may allow the design of direct microbial interventions and open new avenues for preventive measures for breast cancer.

Here we investigated the impact of the dietary botanicals, BSp or GTPs and their combination, on the gut microbiota of Her2/neu mice when these compounds were administered lifelong from conception and from the beginning of early life. We evaluated the impact of these dietary treatments on ER (-) mammary cancer prevention in these mice by assessing the tumor volume percentage and average tumor latency. Gut microbial communities are known for direct interaction with the host as well as indirect interactions via production of diverse metabolites in the host [43]. In this study, we performed 16S rRNA gene sequencing to investigate the gut microbial composition of Her2/neu mice before and after tumor onset, and identified key bacterial phylotypes that were significantly altered with dietary treatment. We also studied the impact of BSp, GTPs and the combination diet on the plasma levels of SCFAs, which are gut microbial-produced metabolites.

## Materials and Methods

### Animals

The animal study was reviewed and approved by Institutional Animal Use and Care Committee of the University of Alabama at Birmingham (IACUC; Animal Project Numbers: 10088 and 20653). The Wild type (WT) Her2/neu [FVB-Tg(MMTV-Erbb2)NK1Mul/J] mouse model (Jackson Laboratory, Bar Harbor, ME) was used in this study. We obtained the breeder mice (4 wks) that were bred from 10 wks of age to obtain sufficient colonies for follow-up experiments. We performed a standard PCR analysis with tail DNA of mice (3 wks of age) to identify the Tag genotypes [44]. Mice were housed in the Animal Resource Facility at the University of Alabama at Birmingham and were maintained within 12-hour light/dark cycle, 24 ± 2°C temperatures, and 50 ± 10% humidity. An online Power and Sample Size Calculator (http://powerandsamplesize.com) was used to evaluate the power and sample size by 2-proportion comparison [26]. All animals had free access to food and water.

### Mouse Diet

Mice on the BSp diet were fed a customized AIN-93G diet from TestDiet (St. Louis, MO) and adjusted for nutrients content as used previously [26]. The modified AIN-93G diet contained 26% (w/w) BSp, which was obtained from Natural Sprout Company (Springfield, MO). Mice on GTPs diet were orally fed 0.5% (w/v) GTPs Sunphenon 90D (SP90D, Taiyo Inc., Minneapolis, MN, USA) in drinking water either alone or in combination with the 26% BSp. The SP90D contained polyphenols (>90%), catechins (>80%), EGCG (>45%) and caffeine (<1%), and is a decaffeinated extract of green tea containing purified polyphenols rich in green tea catechins. Mice on control diet were fed with AIN-93G basal mix diet pellets. Diets were stored in airtight containers and were kept away from light under refrigeration (2°C for up to six months and −20°C for long-term life) to provide maximum protection against possible changes. Mice food was in the form of pellets.

### Animal Experiment Design

#### Beginning in early life (BE) group

80 Her2/neu female mice were randomly divided into four dietary treatment groups (20 mice/group) as the following: control or BSp or GTPs or combination, upon weaning. The dietary treatment continued from 3 wks of age (prepubescence) through adulthood until termination (Fig 1).

**Fig 1.**
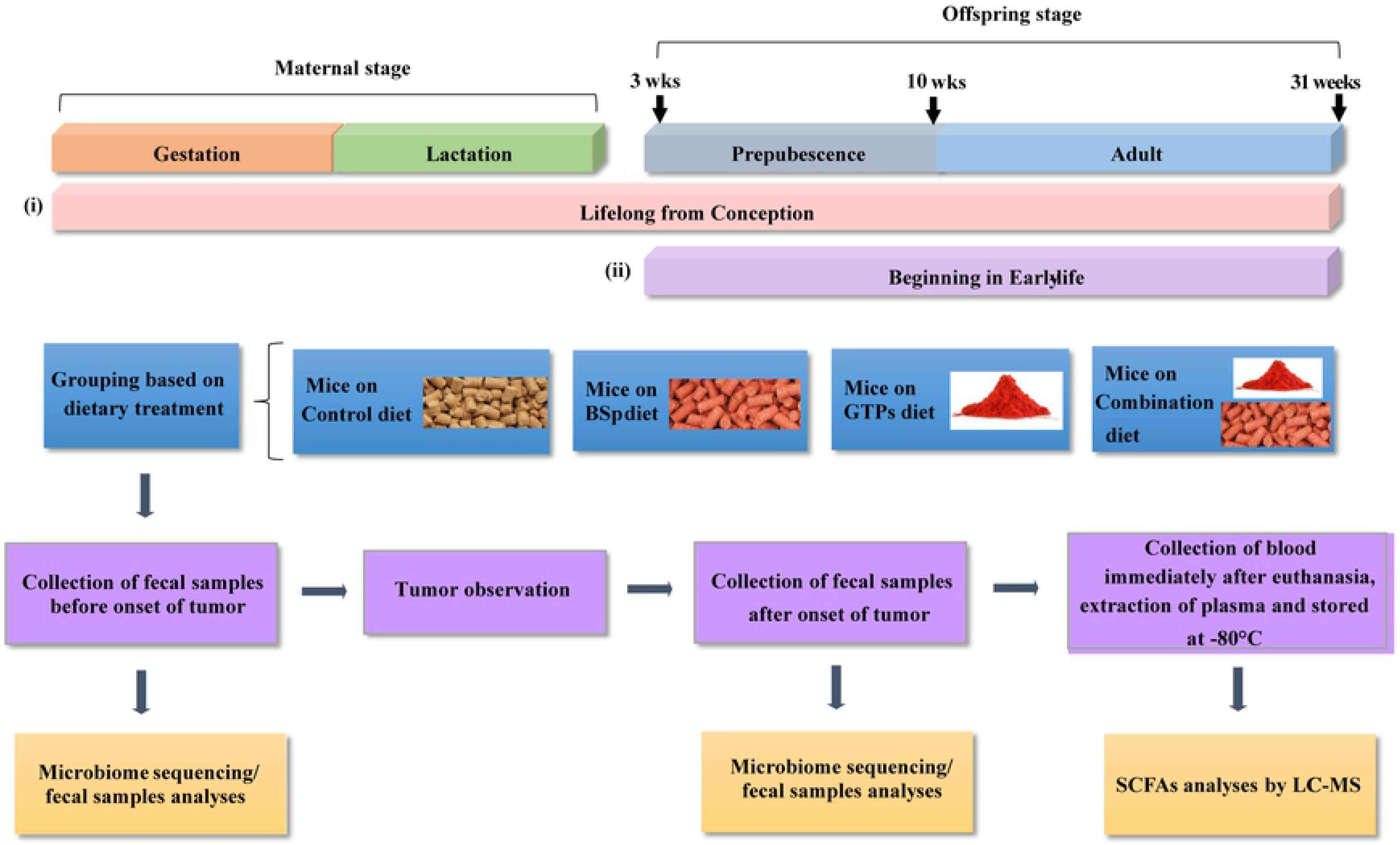
Schematic representation of study design. Her2/neu transgenic mice were administrated either the control diet or 26% broccoli sprouts (BSp) diet in pellets or 0.5% green tea polyphenol (GTP) in the drinking water or BSp and GTPs in the combination diets at two time points: (i) Lifelong from conception (LC), and (ii) Beginning from early life (BE). In the LC group, dietary treatments were started upon the mating of maternal mice and continued during gestation and lactation. After weaning, offspring female mice selected from each group were maintained on the same treatments as their mother throughout their lifespan until termination of the experiment and monitored for tumor growth weekly. In the BE group, female mice were fed one of four different dietary regimens upon weaning, which continued throughout the study until termination. Fecal samples were obtained before the onset of tumor (at 16 wks of age) for the analyses of intestinal communities composition with microbiome analyses. Prior to euthanasia, the fecal samples were obtained after the onset of tumor (at 28 wks of age) for investigation of temporal changes in microbial composition by microbiome analyses. On the day of euthanization, blood samples (approximately 500 μl in Eppendorf tubes containing EDTA) were individually collected from the retroorbital sinus and plasma was isolated by centrifugation, and stored at −80°C for SCFAs analysis.

#### Lifelong from conception (LC) group

40 Her2/neu female mice were randomly divided into four dietary treatments groups (10 mice/group) as the following: control or BSp or GTPs or combination upon pregnancy. The dietary treatments were continued throughout the gestation and lactation periods. After the lactation period, their female offspring mice were weaned at 3 wks of age. Twenty offspring female mice were then randomly selected from each group and fed the same diet as their mother throughout the study until termination.

### Tumor observation and sampling

The tumor size and latency were measured and calculated weekly. Tumor volume was determined as, tumor volume (cm^3^) = 0.523 × [length (cm) × width^2^ (cm^2^)][45]. The experiments were terminated when all of the control mice developed tumors and had an average tumor diameter exceeding 1.0 cm. On the day of euthanization, blood samples (approximately 500 μl in Eppendorf tubes containing EDTA) were individually collected from the retroorbital sinus of each mouse, and plasma was isolated by centrifugation before storing at −80°C for SCFAs analysis. All animal studies were carried out in accordance with the guidelines of the IACUC at UAB.

### Fecal sample collection

The mammary tumors originate at around 20 wks of age in the Her2/neu transgenic mice model [46]. Therefore, the fecal samples from mice were collected at two time points: before the onset of tumor (at 16 wks) and after the onset of tumor (at 28 wks) for studying the temporal efficacy of dietary botanicals on the microbiota composition and impact on breast cancer prevention. Fecal samples were obtained from eight mice per treatment from the BE group and five mice per treatment from the LC group. Approximately 40-50 mg of fecal specimens were acquired and diluted in modified Cary Blair [47] medium for a total volume of 800 μL with 10% by volume glycerol, mixed uniformly by vortex and stored at −80°C [47].

### DNA Extraction and PCR amplification

Genomic DNA was isolated from fecal samples by bead-beating with the Fecal DNA Isolation Kit from Zymo Research (Irvine, CA, USA) according to the manufacturer’s instructions. The extracted DNA was immediately used for PCR or stored in standard Tris-EDTA buffer (pH 8) at 4°C. Before PCR, the isolated PCR DNA was quantified using a microspectrophotometer (ThermoFisher, Waltham, MA) [47]. An amplicon library was constructed from isolated DNA samples by PCR to amplify the V4 region of the 16S rRNA gene with the unique barcoded primers [32], and the olignonucleotide primers were as follows (Eurofind Genomics, Inc., Huntsville, AL):

Forward V4:

5’AATGATACGGCGACCACCGAGATCTACACTATGGTAATTGTGTGCCAGCMGCCGC GGTAA-3’;

Reverse V4:

5’CAAGAGAAGACGGCATACGAGATNNNNNNAGTCAGTCAGCCGGACTACHVGGGT WTCTAAT-3’

The quantification of purified PCR products was carried out by PICO green dsDNA Reagent.

### Illumina MiSeq sequencing and bioinformatics analyses

Agarose gel electrophoresis was performed on the individual PCR products and visualized on the UV illuminator. The isolated PCR products were excised from the gel and purified by QIAquick Gel Extraction Kit (Qiagen, Germantown, MD). NextGen sequencing Illumina MiSeq [32, 47] platform was used for sequencing the PCR products of about 250 bp paired-end reads from the V4 region of the *16S rRNA* gene. The obtained raw FASTQ files were used for library construction, de-multiplexed, and assessed for quality control using FastQC (FastQ quality control). Subsequently, the processed library was used for downstream analyses using the Quantitative Insight into Microbial Ecology (QIIME) [48] data analysis package. As a result the samples were grouped using Uclust in-built function, a clustering program and the sequences of 97% similarity were grouped into Operational taxonomic units (OTU). The multiple sequence alignment of OTUs was created by using PyNAST [49]. Beta diversity was evaluated with Bray Curtis method to quantify continuous dissimilarity between different treatment groups (control-BSp, control-GTPs and control-combination) [50].

### LC-MS analysis of plasma short chain fatty acids

Plasma samples (20 μl) were mixed with ice-cold methanol (60 μl) to precipitate proteins. The methanol contained the internal standard ^13^C_4_-butyric acid (0.5 mg/ml). The samples were centrifuged for 10 min at 16,000 x g and supernatants collected. Each supernatant (40 μl) was diluted into 50% methanol (40 μl) in a 1.5 ml microfuge tube. 1-Ethyl-3-(3-dimethylaminopropyl) carbodiimide (10 μl, 0.25 M) and 10 μl of 0.1 M O-benzylhydroxylamine (o-BHA, 10 μl, 0.1 M) were added to samples to chemically modify SCFAs with o-BHA. The resulting mixture was derivatized for 1 h at room temperature. Samples were diluted 20-fold in 50% methanol. The diluted samples (200 μl) were subject to liquid-liquid extraction with dichloromethane (DCM, 600 μl). Samples were vortexed for 1 min and phases were allowed to separate. A portion of the DCM phase (400 μl) was transferred to a glass tube and dried under N_2_ gas [51]. Samples were reconstituted in 30% methanol (200 μl) and then transferred to loading vials. SCFA standards (MilliporSigma, CRM46957, Burlington, MA) were processed in the same manner as samples. Concentrations from 0.1 – 5,000 μM were used to create standard curves for each SCFA.

Samples were analyzed by tandem HPLC-MS utilizing a 20A HPLC (Shimadzu, Kyoto, Japan) and an API 4000 triple quadrupole mass spectrophotometer (SCIEX, Framingham, MD). Instrument control and data acquisition utilized Analyst 1.6.2 (SCIEX). Authentic standards and samples were analyzed as previously described [51] with slight alterations. An Accucore C_18_ reverse-phase column (2.6 μm 100 x 2.1 mm ID, ThermoFisher, Waltham, MA) was employed for gradient separation. Mobile phase B was altered to 20% isopropanol/80 methanol/0.1% formic acid. MultiQuant 1.3.2 (SCIEX) was used for post-acquisition data analysis; peaks in all standards and plasma extracts were normalized to the ^13^C_4_-butyric acid internal standard signal. Each standard curve was regressed linearly with 1/x^2^ weighting.

### Statistical analysis

Power calculations for animal experiments were conducted using an online calculator (http://powerandsamplesize.com/). Sample size for animal studies was calculated by one-side 2-propotion comparison. Tumor growth was calculated by using Bonferroni adjustment for multiple comparisons between the dietary treatment groups (80% power, significance level of 0.01, alpha = 0.05 with Bonferroni adjustment for 4 comparisons). Statistical analysis of tumor growth data was performed by SPSS version 24.0. The comparisons between two groups were analyzed by two-tailed Student’s *t*-test and comparisons between three or more groups were analyzed by one-way independent ANOVA, followed by Tukey’s post-hoc test to determine significance between groups for tumor volume and tumor latency. The statistical significance of bacterial abundance at the taxonomic level was accounted for by adjusting the false-discovery rate (FDR) at 5% by the Benjamini and Hochberg step-up method [52]. Furthermore, to visualize the correlation between microbiome data (OTUs) among different samples, heatmaps were constructed using pheatmap package in R (v3.6.0). The heatmaps were generated based on Pearson correlation coefficient (R), wherein the rows represent different treatment groups (control, BSp, GTPs and combination) and columns represent different bacterial taxa. Heatmaps of the bacterial taxonomic distribution were generated based on the relative abundance of each bacterial taxa in each dietary treatment for both BE and LC groups. Error bars of tumor growth data were standard error of the mean obtained from experiments. Values of SCFAs data are represented as mean ± SD. Statistically significant results were represented as ** (*p* < 0.01) and * (*p* < 0.05).

## Results

### Effects of the BSp, GTPs or combination dietary treatment on ER-negative mammary tumor development

The Her2/neu female transgenic mouse model is an excellent preclinical model for breast cancer prevention studies because the mice develop spontaneous ER-negative mammary cancer that resembles human pathogenesis [53, 54]. The mice develop focal hyperplastic and dysplastic mammary tumors (due to the overexpression of the *Her2/neu* gene) at an early age (~20 wks) [46]. Fig 2 shows the differences in tumor growth volume and percentage over the whole population, and mean tumor latency between our dietary treatment groups. The mice on BSp or GTPs dietary treatment showed suppression in tumor growth and the combination dietary treatment rendered the strongest inhibiting effect on tumor growth volume in both BE (Fig 2a) and LC (Fig 2b) groups. For all three dietary treatments, the tumor latency was significantly increased in both BE (*p* < 0.01, Fig 2c) and LC (*p* < 0.01, Fig 2d) groups. Therefore, the combination diet group was the most effective in suppressing the tumor development and the BSp diet group was integrally more efficacious than the GTPs group when compared to control.

**Fig 2.**
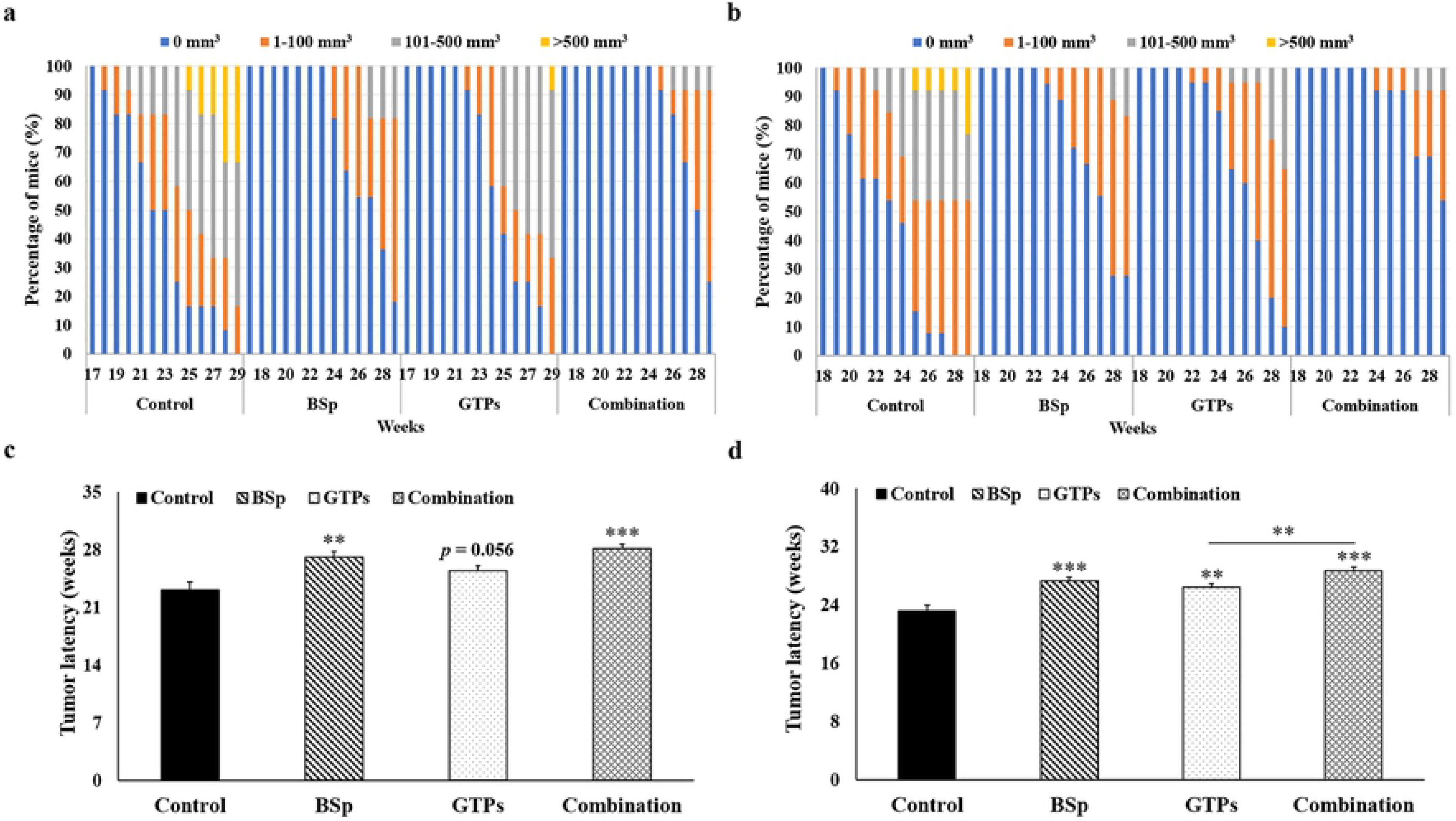
Tumor growth comparisons between control, BSp, GTPs and combination-fed mice dietary groups. Female Her2/neu mice were monitored for tumor growth weekly. (a) The suppression in tumor growth volume and the percentage over the whole population in BSp, GTPs and combination dietary regimens as compared to control in the BE group. Colors indicate different tumor volume ranges as shown in the legend above the graph. (b) The inhibition of tumor growth volume in BSp-fed, GTPs-fed and combination-fed mice when compared to control-fed in the LC group. (c) The mean tumor latency for four different treatments in the BE group. (d) The mean tumor latency of mice in BSp, GTPs, combination and control treatment in the LC group. Columns, mean; Bars, standard error; *, *p* < 0.05; **, *p* < 0.01, ***, *p* < 0.001, significantly different from the control group.

### Effects of the BSp, GTPs or combination diet on gut bacterial diversity before the onset of tumors

These Her2/neu female transgenic mice begin to develop ER(-) mammary tumor at around 20 wks of age [46]. Therefore, we chose the 16^th^ week of age for collection of fecal samples as an initial time point to investigate the effects induced by BSp, GTPs and combination treatment groups on gut microbiota of Her2/neu female mice before the onset of tumor in both BE and LC groups for temporal analyses. In order to identify the outliers in samples of different treatment groups, a sample dendrogram was generated by performing hierarchical clustering using hclust package in R (v3.6.0) (S1 Fig). As a result, in the BE group, there were no outliers and all the samples were included in further analysis. Subsequently, a 3D Principal Coordinates Analysis (PCoA) plot distance metric (Bray Curtis) was generated. Distinct clustering of the BSp-fed and combination-fed treatment groups as compared to the GTPs-fed and control-fed dietary mice in the BE group was observed (Fig 3a), whereas, the GTPs and control dietary treatments were found to be clustered with each other. The results were validated by Permutation Multivariate Analysis of Variance (PERMANOVA) from the distance matrix of the Bray Curtis test of beta diversity in BSp-fed (F = 27.46, *p* = 0.01), GTPs (F = 1.26, *p* = 0.25), and combination-fed (F = 27.08, *p* = 0.01) when compared with control-fed mice. Therefore, these findings indicate the microbiota of BSp-fed mice and combination-fed mice were different from the microbiota of GTPs-fed mice and control-fed mice, which in turn were strongly similar to each other.

**Fig 3.**
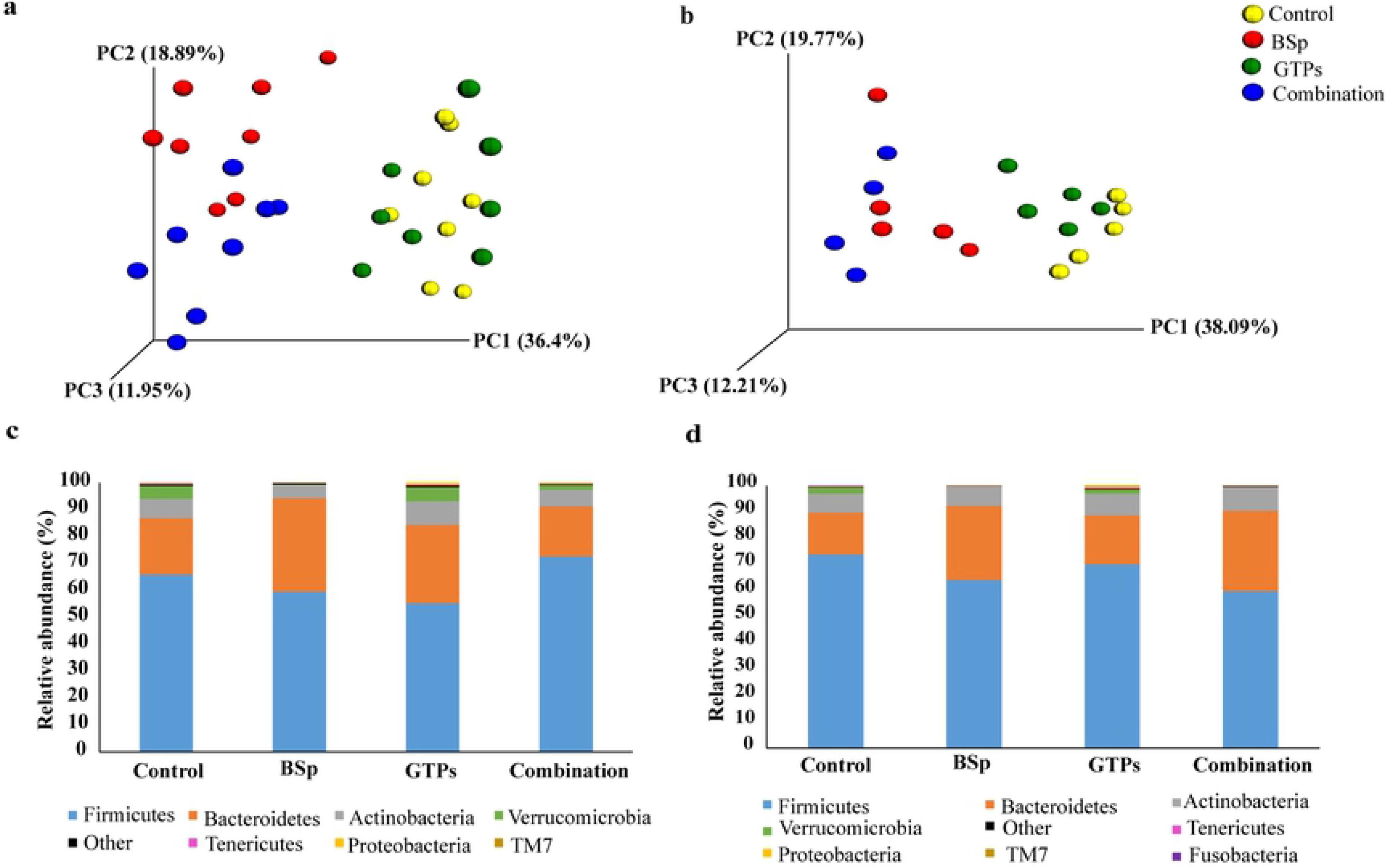
Taxonomic distribution of microbial communities in the gut of mice before the onset of tumor. (a) 3D PCoA plot (Bray Curtis) showing a distinct clustering of the BSp-fed (red) and combination-fed diet groups (blue) as compared to the control-fed (yellow) and GTPs-fed (pink) in the BE group. (b) 3D PCoA plot (Bray Curtis) showing a distinct clustering of the control-fed (yellow), BSp-fed (red), GTPs-fed (pink) and combination-fed (blue) diet groups in the LC group. (c) Pre-tumor phylum level changes in microbial abundance by our dietary treatments in the BE group. (d) Pre-tumor phylum level changes in microbial composition by our dietary treatments in the LC group.

Similarly, hierarchical clustering using hclust package in R (v3.6.0) in the LC temporal treatment group was performed and no outliers were identified (S2 Fig); thus, we included each sample in this study. We observed an overall similar clustering as seen in the BE group, the microbial composition of BSp-fed mice strongly overlapped with combination-fed mice and the microbial composition of GTPs-fed mice strongly overlapped with control-fed mice (Fig 3b). This observation was validated with PERMANOVA from the distance matrix generated using Bray Curtis method with beta diversity. It was significant in BSp-fed (F = 12.69, *p* = 0.02) and combination-fed (F = 37.61, *p* = 0.03) mice when compared with control-fed mice. The microbial composition of GTPs-fed was highly clustered but it failed to show a clear separation from the microbial composition of control-fed (F = 2.78, *p* = 0.08) mice. Hence, these results suggest that the microbial composition of BSp-fed mice and combination-fed mice were largely similar and the microbial composition of GTPs-fed mice and control-fed mice were lately similar to each other.

### Impact of diet on gut bacterial composition before the onset of tumor

When compared at the phylum level, the dietary treatments showed differences in microbial abundances in both the BE and LC groups. In the BE group, the mice that were BSp-fed underwent a significant increase in Bacteroidetes as compared to control-fed mice (35% versus 21%, *p* = 0.02) (Fig. 3c). The microbiota of mice on the GTPs diet showed a significant decrease in Firmicutes phylum (55% versus 66%, *p* = 0.031) as compared to control-fed mice. The microbiota of mice on the combination dietary treatment underwent a significant decline in the abundance of Verrucomicrobia as compared to the mice fed with the control diet (2% versus 5%, *p* = 0.03). The relative abundance of the top 25 bacterial taxonomic units are depicted in a heat map with clustering dendrogram (Fig 4a, S1 Table). Compared to the control diet, the relative abundance of Firmicutes (f_Erysipelotrichaceae g_*Allobaculum* s_unclassified) was higher in the BSp-fed and combination-fed mice. Family Bifidobacteriaceae and Lactobacillaceae showed higher abundance in the control-fed and GTPs-fed when compared to BSp-fed and combination-fed diets. After investigation of significance of bacterial communities using false-discovery rate (FDR) correction, numerous significantly different bacterial taxa (S1 File) were found. Table 1 shows the top bacterial taxa that were different between the BSp-fed, GTPs-fed and combination-fed as compared to control-fed mice groups, respectively. *Allobaculum* levels were found to be significantly increased in the BSp-fed (5.5-fold) group and in the combination-fed (8.3-fold) group. The S24-7, Ruminococcaceae family and *Lactococcus* were increased significantly after consumption of the BSp diet and combination diet in mice. Mice on GTPs dietary treatment did not show any significant changes in bacterial communities after the correction for multiple comparisons (FDR < 0.05). Overall, the BSp-fed and combination-fed groups showed significant changes in the microbial abundance as compared to the control diet at the taxonomic level.

**Table 1.**
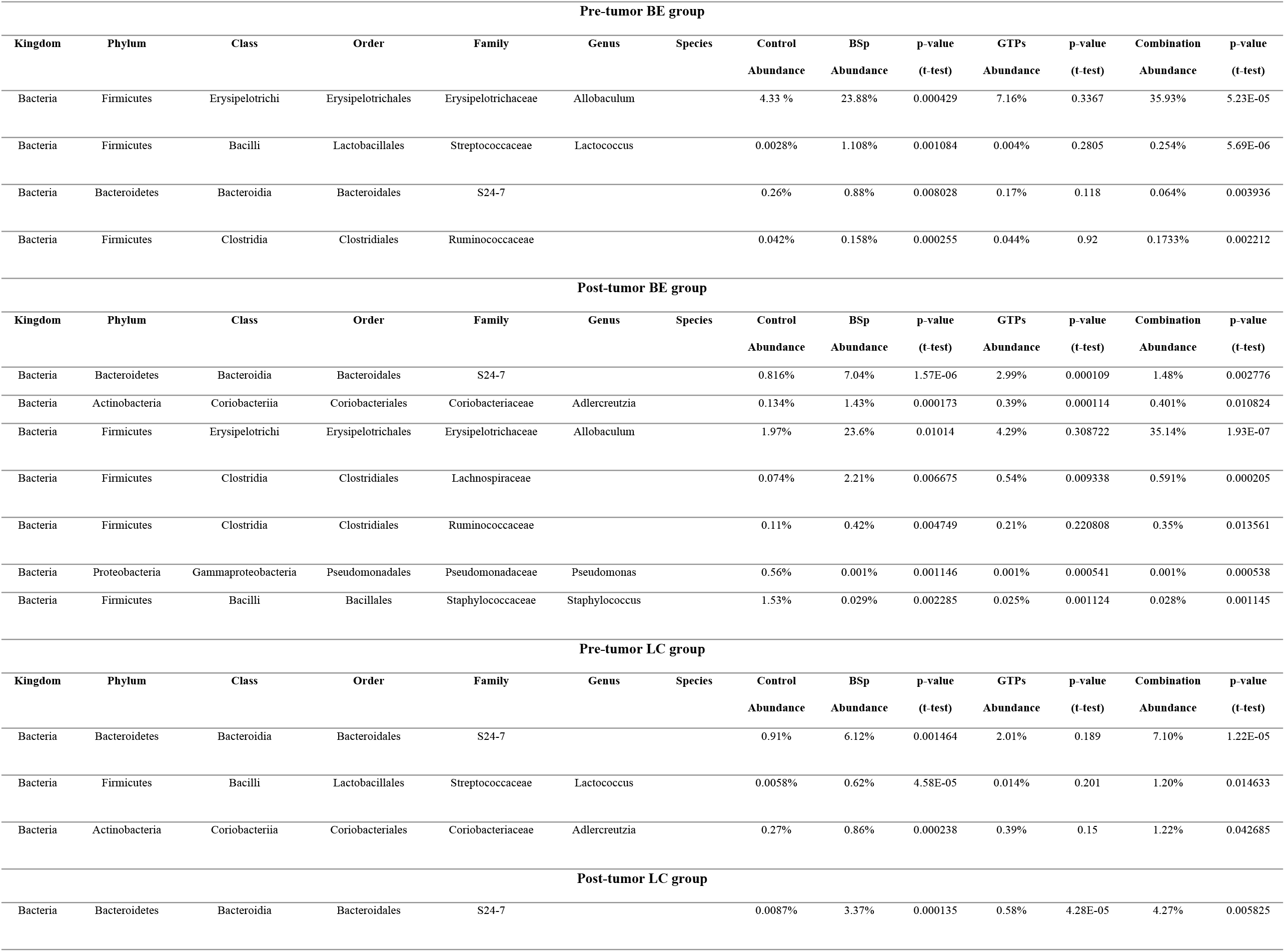
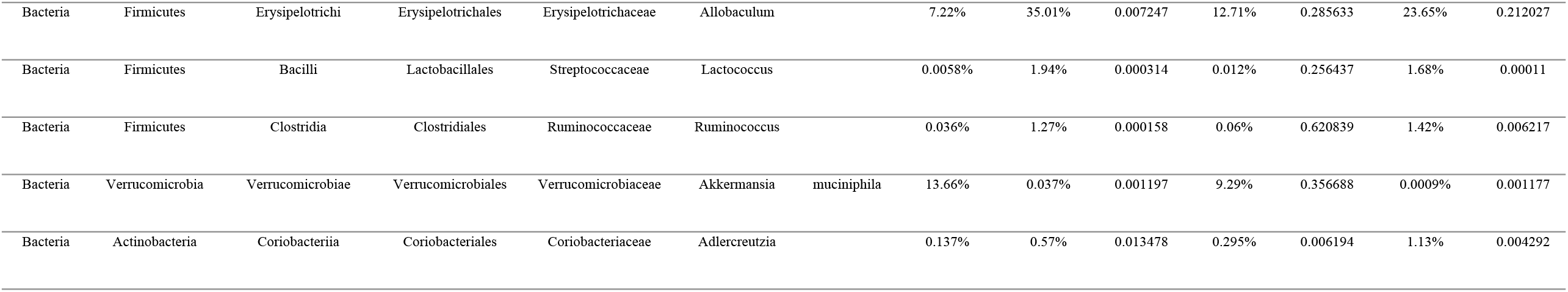
The significant changes in bacterial species relative abundance after the onset of tumor in BSp-fed, GTPs-fed and combination-fed mice versus control-fed mice of both BE group and LC group.

**Fig 4.**
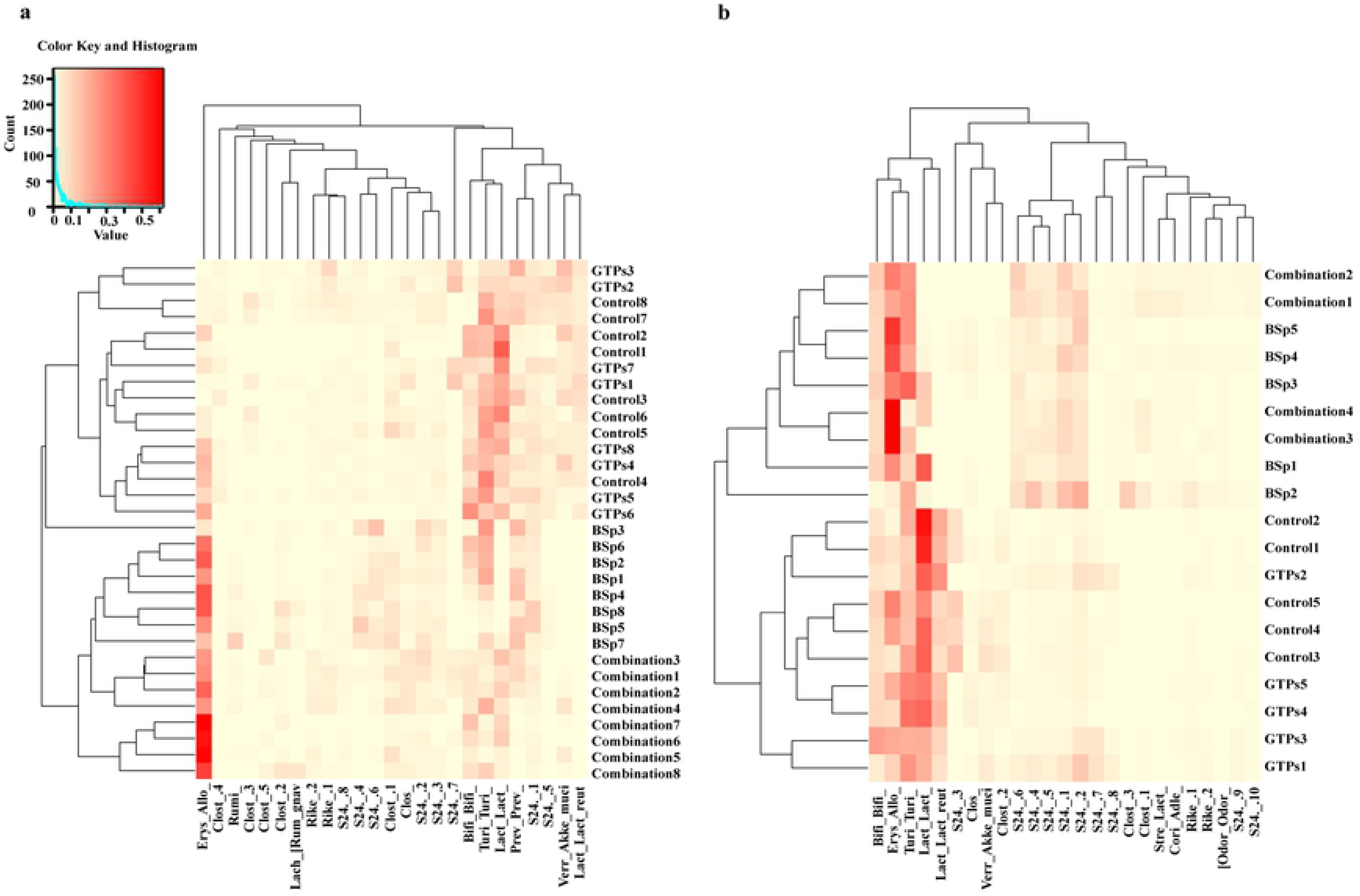
Heatmaps showing the relative abundance of microbial species in mice before the onset of tumor. (a) Heatmap of the bacterial abundance between BSp-fed, GTPs-fed, combination-fed and control-fed mice before the onset of tumor in the BE group. (b) Heatmap of the microbial species abundance between BSp, GTPs, combination and control dietary treatment before the onset of tumor in the LC group. The dendrogram shows the distinct clustering of BSp and combination dietary treatments and the distinct clustering of the GTPs and control diet groups. Red color depicts higher abundance; yellow color depicts lower abundance. The lineages are abbreviated and their details have been provided in S1 and S2 Tables.

At the phylum level, the mice in the LC group showed a similar trend as the mice in the BE group. Mice on the BSp diet (28% versus 16%, *p* = 0.04) and combination diet (31% versus 16%, *p* = 0.005) underwent a significant increase in Bacteroidetes levels as compared to mice fed the control diet (Fig 3d). The relative abundance of the top 25 bacterial taxonomic units are depicted in the heatmap with clustering dendrogram (Fig 4b, S2 Table). Similar to the BE group, the relative abundance of Firmicutes (f_Erysipelotrichaceae g_*Allobaculum* s_unclassified) was higher in the BSp-fed and the combination-fed mice. After correction for multiple comparisons (FDR < 0.05), various microbial communities were found to be significantly different (S1 File) in the dietary treatment groups. The top bacterial species that were different in the mice on BSp, GTPs and combination dietary treatments and control are shown in Table 1. The S24-7 family, *Lactococcus* and *Adlercreutzia* bacteria was increased significantly after the consumption of the BSp or combination diet. The microbiota of GTPs-fed mice did not show significant increase in any bacterial family after the correction for multiple comparisons (FDR < 0.05) when compared with the control-fed mice. Overall, the mice on BSp and combination treatment groups displayed significant differences in bacterial species as compared with the control group of mice.

### Effects of the BSp, GTPs or combination diet on gut bacterial diversity after the onset of tumor

We chose another time point (28^th^ wks) for collection of fecal samples to investigate whether the gut microbiota was altered after the onset of tumor by our dietary treatments in mice. We sought to investigate changes in gut microbial composition with 3D PCoA plot distance metric (Bray Curtis) and found a distinct clustering of microbial communities with the treatment of BSp, GTPs or combination as compared to the control diet in the BE group (Fig 5a). PERMANOVA test on the Bray Curtis clustering supported our observation: BSp (F = 28.88, *p* = 0.01), GTPs (F = 11.16, *p* = 0.01) and combination (F = 35.10, *p* = 0.01) dietary treatments demonstrated significant clustering against the control group. In addition, the microbiota of BSp-fed and combination-fed mice were clustered together again, which implied a similar distribution of bacterial species even after the tumor onset.

**Fig 5.**
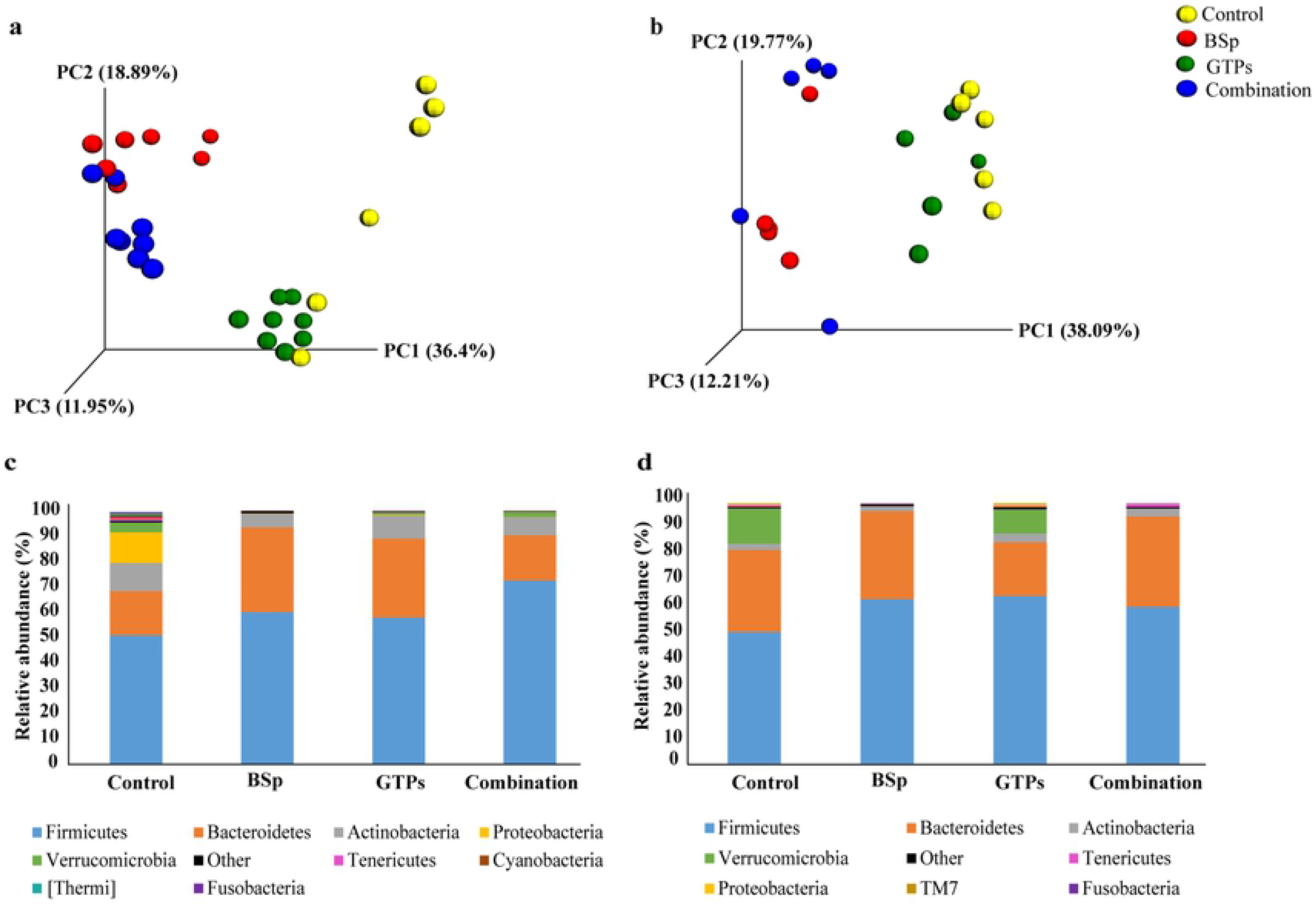
Changes in microbial composition by the dietary treatments after the onset of tumor in mice. (a) After tumor onset, the 3D PCoA plot (Bray Curtis) showed a distinct clustering of the BSp-fed (red), GTPs-fed (pink) and combination-fed diet groups (blue) as compared to the control-fed (yellow) in the BE group. (b) 3D PCoA plot (Bray Curtis) showed a distinct clustering of dietary groups after the tumor onset in the LC group. (c) Post-tumor phylum level changes in microbial abundance by our dietary treatments in the BE group. (d) Post-tumor phylum level changes in microbial abundance by our dietary treatments in the LC group.

In the LC group, we observed distinct clustering of the mice on BSp diet or GTPs diet or combination diet or control diet (Fig 4b). PERMANOVA test on the Bray Curtis clustering validated our findings-BSp (F = 11.43, *p* = 0.01), GTPs (F = 3.25, *p* = 0.03) and combination (F = 11.43, *p* = 0.01). The same microbial clustering pattern of BSp with combination, and GTPs with control is observed as was reported in pre-tumor evaluation.

### Impact of diet on gut bacterial composition after onset of tumor

The changes in phylum levels of microbial composition were also observed after tumor onset for all dietary treatments of both groups. In the BE group, the mice on the BSp diet showed a significant increase in Bacteroidetes (33% versus 18%, *p* < 0.01) and a significant decrease in Actinobacteria (6% versus 11%, *p* = 0.02) and Proteobacteria (0.14% versus 12%, *p* < 0.01) as compared to mice on the control diet (Fig. 5c). The mice on GTPs diet had a significant increase in the Bacteroidetes levels (31% versus 18%, *p* < 0.001) and a significant decrease in Proteobacteria (0.28% versus 12%, *p* < 0.01) as compared to mice on the control diet. The mice on the combination dietary treatment revealed a significant increase in Firmicutes (73% versus 51%, *p* < 0.001), and a significant decrease in Proteobacteria (0.11% versus 12%, *p* < 0.01) abundance. The heat map (Fig 6a, S3 Table) depicts the abundance of family Erysipelotrichaceae (f_Erysipelotrichaceae g_*Allobaculum* s_unclassified) to be high in BSp, GTPs and combination dietary treatments. In addition, the *Lactobacillus* showed higher abundance in the GTPs-fed and combination-fed groups when compared to BSp-fed and control-fed groups. We found several significantly different bacterial taxonomic abundance following correction with FDR (<0.05) (S1 File). Table 1 lists the top taxa that had different abundance between the dietary treatments’ post-tumor temporal period. There was a significant increase in *Allobaculum* levels in BSp-fed (~11.979 fold) and combination-fed (~17.837 fold) mice groups when compared to the control-fed mice. The S24-7, Lachnospiraceae family of bacteria and *Adlercreutzia* genus were increased significantly in mice of all dietary treatments when compared with the control treatment. Specifically, *Pseudomonas* and *Staphylococcus* genus levels were significantly lowered in all dietary treatments administered to mice. Collectively, an overlap of numerous bacterial communities was observed due to the dietary treatments that were significantly different from microbial communities of the control group mice.

**Fig 6.**
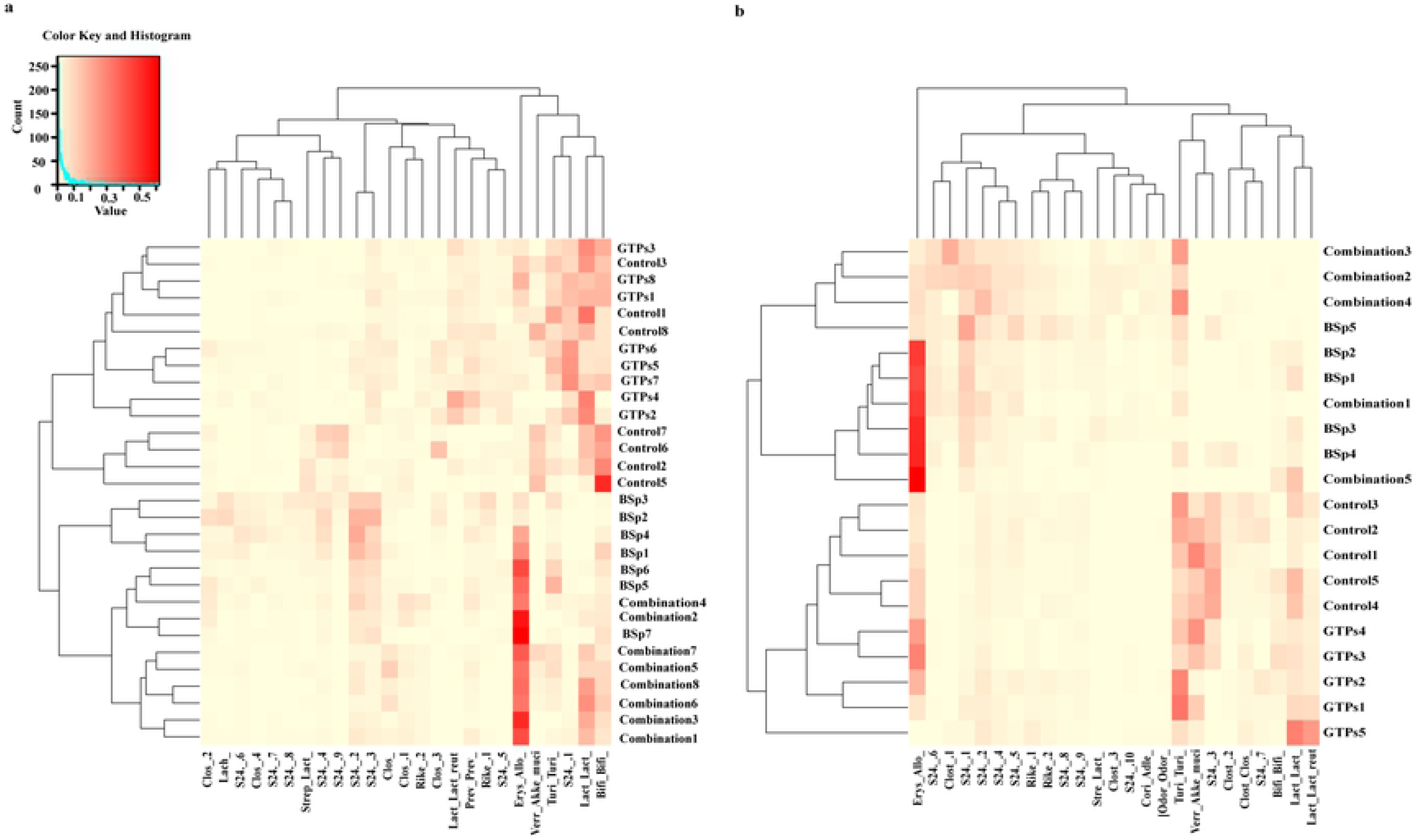
Heatmaps showing the relative abundance of microbial species after the onset of tumor in mice. (a) Heatmap of the bacterial abundance between BSp-fed, GTPs-fed, combination-fed and control-fed mice after tumor onset in the BE group. (b) Heatmap of the microbial species abundance between dietary treatments after tumor onset in LC group. Dendrogram shows the clustering of BSp and combination dietary treatment with each other and the clustering of the GTPs and control diet groups with each other. Red color depicts higher abundance; yellow color depicts lower abundance. The details of abbreviated lineages have been provided in S1 and S2 Tables.

The alterations in microbial compositions were also observed in the LC group after the onset of tumors (Fig 5d). BSp treatment led to a significant decrease in levels of Proteobacteria (0.07% versus 0.61%, *p* = 0.001). GTPs treatment led to significantly higher levels of Firmicutes (65% versus 51%, *p* = 0.026) and also significantly lower levels of Bacteroidetes (21% versus 31%, *p* < 0.001) in comparison to the control-fed mice. Furthermore, the combination treatment led to a significant decrease in Proteobacteria (0.07% versus 0.61%, *p* = 0.001) when compared with the control-fed mice. The heatmap (Fig 6b) indicated the augmentation of bacteria of genus *Allobaculum* in BSp-fed and combination-fed diet groups when compared to the control group. There were several significantly different bacterial taxa between dietary treatments after correction for multiple comparisons with the FDR correction (<0.05) (S1 File). Table 1 displays the top microbial species that had different abundance between the dietary groups of mice after the onset of tumor. Some of the same bacterial communities have been found to be changed such as family S24-7 and *Adlercreutzia* genus that were significantly increased when compared with the control group. Overall, these results show that dietary treatments led to significant changes in microbial species after the onset of tumors.

### Analyses of the quantity and type of SCFAs in mice on the BSp, GTPs, combination or control diet

Short-chain fatty acids are the crucial metabolites produced from fermentation of dietary fiber by intestinal communities, and act as signaling molecules in the complex crosstalk network of the gut with distal organs [55]. Therefore, we investigated whether the plasma SCFA profiles changed with our dietary treatments. As shown in Fig 7a, we detected the levels of propionate, butyrate, isobutyrate, valerate and hexanoate in the plasma samples of mice fed with BSp diet, GTPs diet, combination diet or control diet. In the BE group, the plasma concentrations of all SCFAs were found to be unchanged in BE-fed, GTPs-fed and combination-fed mice when compared with the plasma levels of control-diet mice.

**Fig 7.**
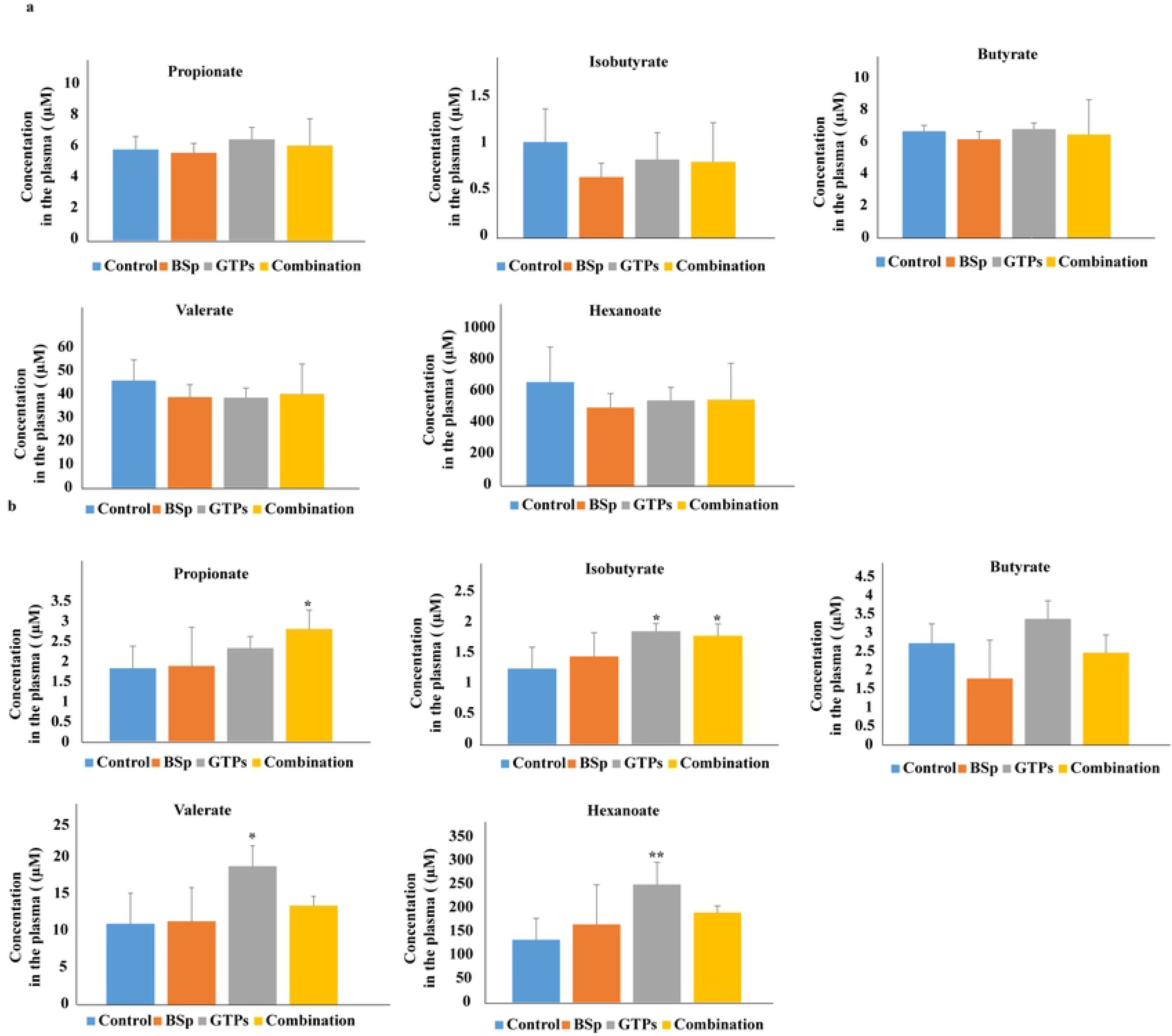
Analyses of plasma SCFAs levels in BSp-fed, GTPs-fed, combination-fed or control-fed mice. (a) Concentrations of SCFAs propionate, isobutyrate, butyrate, valerate and hexanoate in our dietary treatment mice versus control treatment mice in BE group (b) Concentrations of the SCFAs in BSp-fed, GTPs-fed or combination-fed mice when compared with control-fed mice in LC group. Data are presented as mean ± standard deviation (SD) (n = 6). * *p* < 0.05, ** *p* < 0.01, \*\*\**p* <0.001.

In the LC group (Fig 7b), the BSp-fed mice showed a no effect on the levels of SCFAs in comparison to the control-fed mice. Strikingly, however GTPs-fed mice showed a significant increase in isobutyrate (1.5 fold, *p* = 0.017), valerate (1.48 fold, *p* = 0.02) and hexanoate (1.88 fold, *p* < 0.01) as compared to the control-fed mice. The mice on the combination diet had significantly higher levels of propionate (1.5 fold, *p* = 0.037) and isobutyrate (1.44 fold, *p* = 0.036) in contrast to mice on the control diet. Therefore, the mice on GTPs and combination diet groups had significantly increased levels of SCFAs in the LC group.

## Discussion

The use of dietary bioactive compounds as an adjuvant therapy for chemoprevention of breast cancer has been of great interest. The gut microbiome is an important determinant of human health [56] and alterations in composition and diversity of colonizing microbial communities have been associated with pathogenesis of a vast number of disorders over the past decade [57–60]. We investigated the impact of early life consumption of these dietary compounds on the inhibition of breast cancer, intestinal communities and their derived metabolites in Her2/neu mice by analyzing the gut microbiota and plasma SCFAs profiles.

Our study involved administration of several diets - the BSp diet, the GTPs diet and their combination diet at two exposure periods, beginning in early life (BE) and lifelong from conception (LC) in the Her2/neu transgenic spontaneous ER (-) mammary cancer mouse model. We found in both treatment plans the combination group achieved maximum efficacy in suppression of mammary tumor volume and significantly delayed the tumor latency, followed by the BSp dietary treatment that was more efficacious than the GTPs group when compared to the control group.

The pre-tumor beta diversity analyses revealed that microbial communities of BSp-fed mice and combination-fed mice clustered with each other, whereas microbial communities of GTPs-fed mice and control-fed mice clustered with each other in both BE and LC groups. This may imply BSp treatment contributes more towards the combination treatment as compared to the GTPs group. To investigate changes induced in the gut microbiota by our dietary treatments, we observed the phylum and taxonomic abundance of bacterial communities residing in the gut of mice. At the phylum level, we found an increase of Bacteroidetes in BSp-fed and combination-fed group in both BE and LC groups. Bacteroidetes phyla is known for SCFAs production and was found to be associated with fiber consumption in fecal samples obtained from the monozygotic twins [61].

At the taxonomic level, an increase of genera *Allobaculum, Lactococcus*, Ruminococcaceae and S24-7 family in BSp-fed and combination-fed mice were observed in the BE group. Pre-tumor analyses of gut microbiota in the LC group showed significant increases in bacteria of genera *Lactococcus, Adlercreutzia* and family S24-7 in the BSp-fed and combination-fed mice. These findings also support the promising impact of BSp on the combination treatment in transgenic mice and also may imply a strong association of bacterial abundance with our dietary treatments in both BE and LC groups.

A study found that an antimicrobial peptide produced by *Lactococcus lactis* showed high cytotoxic effects against the MCF-7 breast cancer cell line [62]. Another study found that exopolysaccharides produced by *L. lactis* enhanced the secretion of tumor necrosis factor alpha and inducible nitric oxide synthase in MCF-7 cells as compared to control cell line in a dosage-dependent manner. In addition, nuclear fragmentation, chromatin condensation, cellular shrinkage and decrease in mitochondrial potential were also observed in treated MCF-7 cells [63]. *Adlercreutzia* is a gram-positive, strict anaerobic bacterium, which has been considered to exhibit beneficial effects and possess immunoregulatory properties [64]. In addition, *Adlercreutzia* is an equol-producing bacterium, which has been isolated from the gut of animals and humans [65]. This bacterium’s levels were reported to be decreased in mice fed with a high fat diet, resulting in lowered S-equol concentration in a prostate cancer transgenic mice model [66].

After the onset of tumor in mice, we sought to investigate the temporal impact induced by our dietary treatment on gut microbiota and found a similar pattern of microbial diversity from beta diversity, as the BSp-fed mice microbial composition was clustered with combination-fed mice and GTPs-fed mice microbiota was clustered with control-fed mice in both BE and LC groups. This observation implies a strong association of the BSp treatment group with the combination group, as compared with the GTPs group. We also found modifications at phylum levels induced by our dietary botanicals. For instance, we observed a significant decrease in Proteobacteria levels in BSp-fed and combination-fed mice of both the BE and the LC groups. Proteobacteria phyla consists of various human pathogens such as *Escherichia*, *Helicobacter*, *Neisseria*, *Salmonella* and *Shigella*, which has been implicated in human diseases [67–69].

The investigation of taxonomic microbial abundance in the BE group revealed a significant increase in *Allobaculum* genus levels in mice on the BSp and combination diets. Furthermore, a significant rise of *Adlercreutzia* genus, S24-7, Ruminococcaceae and Lachnospiraceae families and a significant decrease of *Pseudomonas* and *Staphylococcus* genus levels was observed in all dietary treatment groups versus the control group. Ruminococcaceae are primary butyrate-producing bacterial family members that inhabit a healthy colon [70]. The increase of these bacterial taxa due to our dietary treatments illustrates the profound impact of dietary compounds on the establishment of gut microbiota in transgenic mice.

An increased abundance of the S24-7 family was notable in mice administrated a low-fat diet and, in association with increased exercise, prevented weight gain in C57BL/6 mice [71]. Lachnospiraceae family of bacteria have the ability to generate energy in the host by degradation of polysaccharides in the plants [72]. On the other hand, some *Pseudomonas* species are pathogenic such as *Pseudomonas aeruginosa* which can cause wound infections and cystic fibrosis, and *P. oryzihabitans* which can cause sepsis in humans [73]. Investigation of the effects of heat-killed *P. aeruginosa* and *Staphylococcus epidermidis* found an increase in cellular proliferation of MCF-7 and HT-29 breast cancer cell lines and therefore these bacterial strains acted as cancer-deteriorating agents [73].

In the LC group after the onset of tumor, we found a significant increase in S24-7 family, *Lactococcus* and *Adlercreutzia* genus at taxonomic levels induced by our dietary treatments. In addition, the mice in the LC group showed a significant increase in *Allobaculum* genus levels when on the BSp diet. Studies have reported that *Allobaculum* is a beneficial bacteria that produces SCFAs in the intestine of mice [74]. The *Allobaculum* genus also exhibits diverse functions such as anti-inflammatory processes, protection of intestinal barrier, regulation of immune system and host metabolism [75]. A recent study investigated the impact of probiotics supplement on the gut microbiome in a colitis-associated colon cancer mice model and found a significant abundance in the levels of *Allobaculum* in the probiotic group as compared to the control group of mice [76].

Therefore, our findings may implicit potent impact of our dietary compounds on establishment and composition of gut microbiota pre-tumor and post-tumor of mice. These findings also suggest that consumption of BSp, GTPs and their combination in either the beginning to early life (BE) group or life-long from conception (LC) group results in establishment of almost the same microbial consumption, which implies the strong influence of these dietary compounds and that may be potentially beneficial for health and in cancer prevention.

The relationship between commensal communities residing in the gut and host physiology have been well studied [77]. Certain bacterial communities do not show direct oncogenic effects on the tumor, as they can indirectly lead to tumor inhibition by production of gut-derived metabolites. For instance, studies have indicated that gut microbiota can metabolize lignans present in the edible plants into enterolactone, which can regulate estrogen signaling and may lead to protective effects against breast cancer [78, 79]. We sought to investigate if the potential impact of gut microbiota from our dietary treatments on mammary tumorgenesis is modulated via microbial-produced metabolites. SCFAs, such as acetate, propionate and butyrate are major metabolites produced by gut microbiota [80]. Butyrate is a known HDAC inhibitor [81] and has shown antineoplastic properties in various cancers [82]. Sodium butyrate has shown to induce apoptosis in breast cancer cell lines via production of reactive oxygen species and mitochondrial impairment [83]. Studies have found that propionate and valerate are also HDAC inhibitors [84]. Administration of sodium propionate resulted in inhibition of MCF-7 cellular proliferation in a dose-dependent manner and cell-cycle arrest [85]. Additionally, isobutyrate (a minor SCFA) can exhibit anticarcinogenic effects in colon carcinoma [86]. Hexanoate is a medium chain fatty acid that can contribute to cellular inhibition in colorectal cancer, breast cancer and skin cancer cell lines by down-regulation of cell cycle genes and induction of apoptosis [87]. Moreover, the microbial produced metabolites, such as lithocholic acid (a bile acid) can induce oxidative and nitrosative stress resulting in breast cancer inhibition [88].

## Acknowledgements

The authors would like to thank Dr. Peter Eipers of the Department of Cell, Developmental and Integrative Biology at UAB for assisting with the microbiome analyses. We would also like to thank William Van Der Pol in the Center for Clinical & Translational Science at UAB for assistance with bioinformatics analyses. Appreciation is expressed to Landon S. Wilson from the Targeted Metabolomics and Proteomics Laboratory at UAB for technical assistance.

## Author contributions

TOT and MS designed and conceived the experiments. MS, YL, TFB and SL conducted the experiments. MS, IA, MLS, CM, SB, TFB and TOT analyzed the results. TOT provided the reagents/materials/analysis tools for the study. All authors reviewed and approved the final manuscript.

## List of Abbreviations

BE: Beginning in early life
BSp: Broccoli sprouts
DCM: Dichloromethane
DNMTs: DNA methytransferases
EGCG: Epigallocatechin-3-gallate
ER: Estrogen receptor
FastQC: FastQ quality control
FDR: False-discovery rate
GTPs: Green tea polyphenols
HDAC: Histone deacetylase
HER2: Human epidermal growth factor receptor
LC: Life-long from conception
o-BHA: O-benzylhydroxylamine
OTU: Operational taxonomic unit
PCoA: Principal Coordinates Analysis
PERMANOVA: Permutation Multivariate Analysis of Variance
PR: Progesterone receptor
QIIME: Quantitative Insight into Microbial Ecology
SCFA: Short-chain fatty acid
SFN: Sulforaphane
TNBC: Triple negative breast cancer
WT: Wild type

## Supporting information

**S1 Fig. Sample clustering of BSp, GTPs, combination and control dietary treatment microbial abundance data of the BE group.** This clustering dendrogram shows no outliers and therefore all samples from dietary treatments were used in the fecal samples analyses of BE group.

**S2 Fig. The clustering of microbial abundance data of BSp-fed, GTPs-fed, combination-fed and control-fed mice in the LC group.** This hierarchical clustering dendrogram verifies that no outliers were present and we have therefore included every sample for microbiome analyses of the LC group.

**S1 Table. Naming convention for taxonomic microbial composition before the onset of tumor in the BE group.** The naming convention for the heat map before the onset of tumor in the BE group is depicted in Fig 4a. The kingdom, phylum, class, order, family, genus and species of the top 25 bacterial abundances between the BSp-fed, GTPs-fed, combination-fed and control-fed mice groups are enlisted in this table.

**S2 Table. Naming convention of heat map for taxonomic microbial composition before the onset of tumor in the LC group.** Table shows the naming convention for heat map of LC group as shown in Fig 4b. The table also shows the kingdom, phylum, class, order, family, genus and species of the significantly different bacterial abundances between the BSp-fed, GTPs-fed, combination-fed and control-fed mice groups.

**S3 Table: Naming convention for taxonomic microbial composition after the onset of tumor in the BE group.** The naming convention for the heat map after the onset of tumor in the BE group is depicted in Fig 6a. The kingdom, phylum, class, order, family, genus and species of the top 25 bacterial abundances between the BSp-fed, GTPs-fed, combination-fed and control-fed mice groups are enlisted in this table.

**S4 Table. Naming convention of heat map for taxonomic microbial composition after tumor onset in the LC group.** Table shows the naming convention for heat map of LC group as shown in Fig 6b. The table also shows the kingdom, phylum, class, order, family, genus and species of the significantly different bacterial abundances between the BSp-fed, GTPs-fed, combination-fed and control-fed mice groups.

**S1 File. Changes in relative abundance of bacterial species in BSp-fed, GTPs-fed, combination-fed and control-fed mice in the BE and LC groups.** This sheet enlists the changes in relative abundance of bacterial communities of all treatment groups: BSp-fed, GTPs-fed, combination-fed and control-fed mice, in BE and LC group at both before and after onset of the tumor.

